## Supplemental Figures for "Natural genetic variation underlying the negative effect of elevated CO_2_ on ionome composition in *Arabidopsis thaliana*"

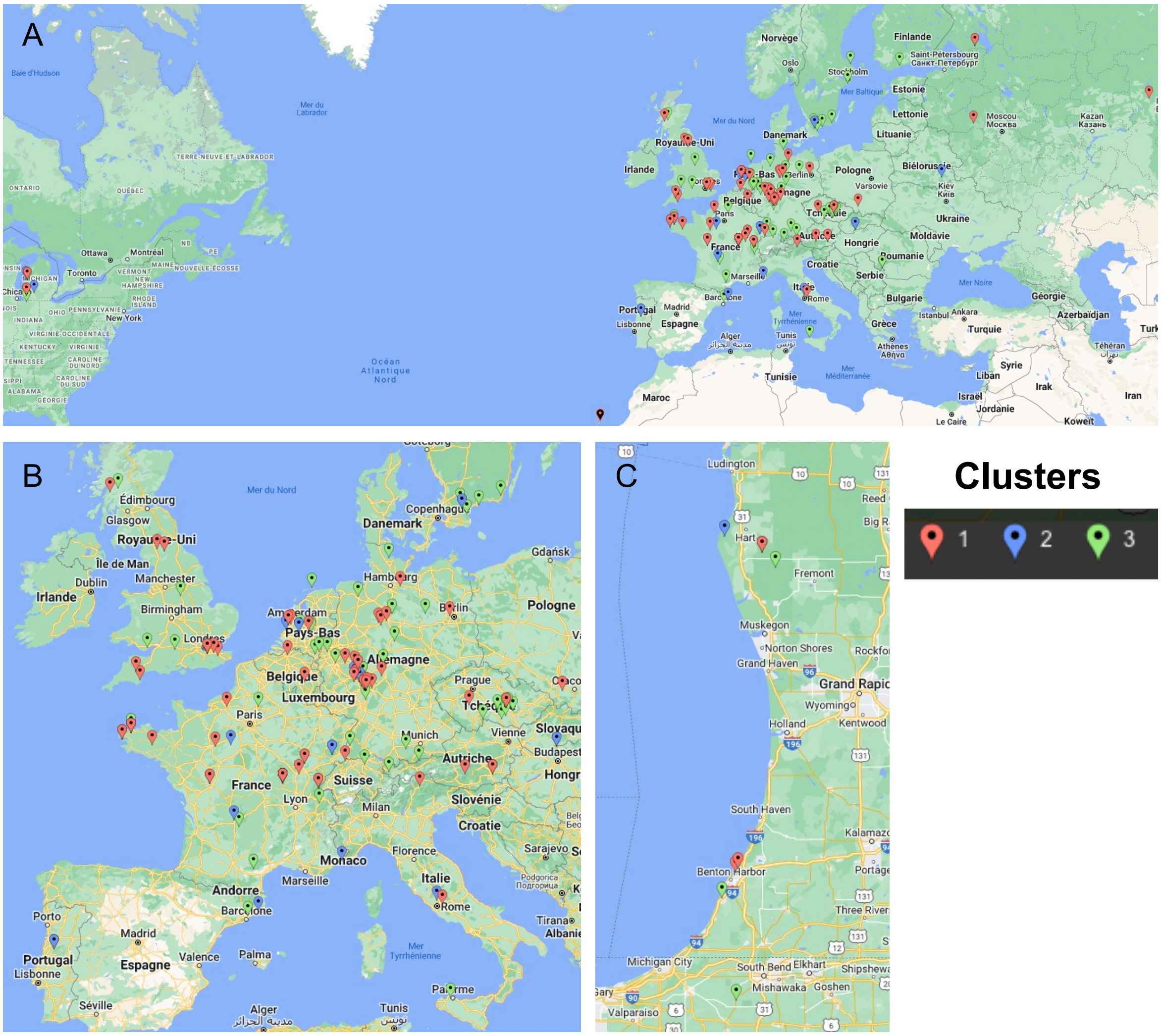


**Supplemental Figure 1:** Geographic distribution of accessions for each cluster identified within the REGMAP panel. A. Full distribution. B. Zoom into European accessions. C. Zoom into North-American accessions.

**A**

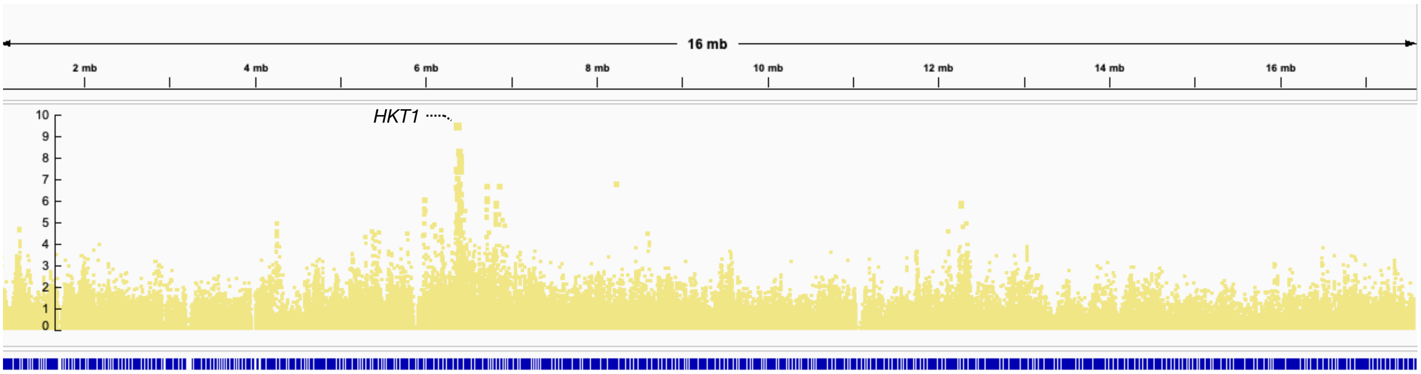


**B**

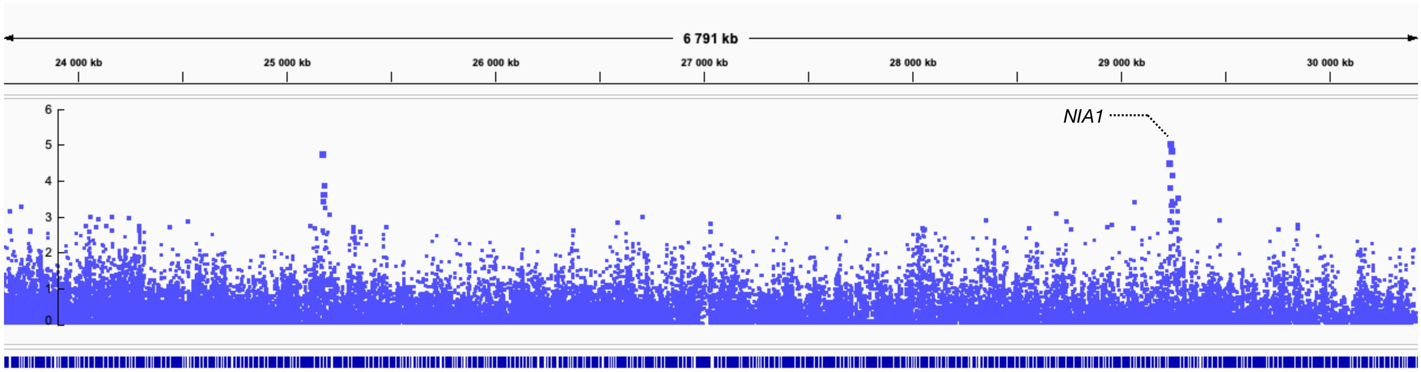


**Supplemental Figure 2:** Snapshots of the Manhattan plots of Na content (**A**) and N content (**B**) under ambient $CO_2$ in the REGMAP panel. Peaks of high P-values are observed at the *HKT1* locus for the Na content (**A**) and at the *NIA1* locus for the N content (**B**).

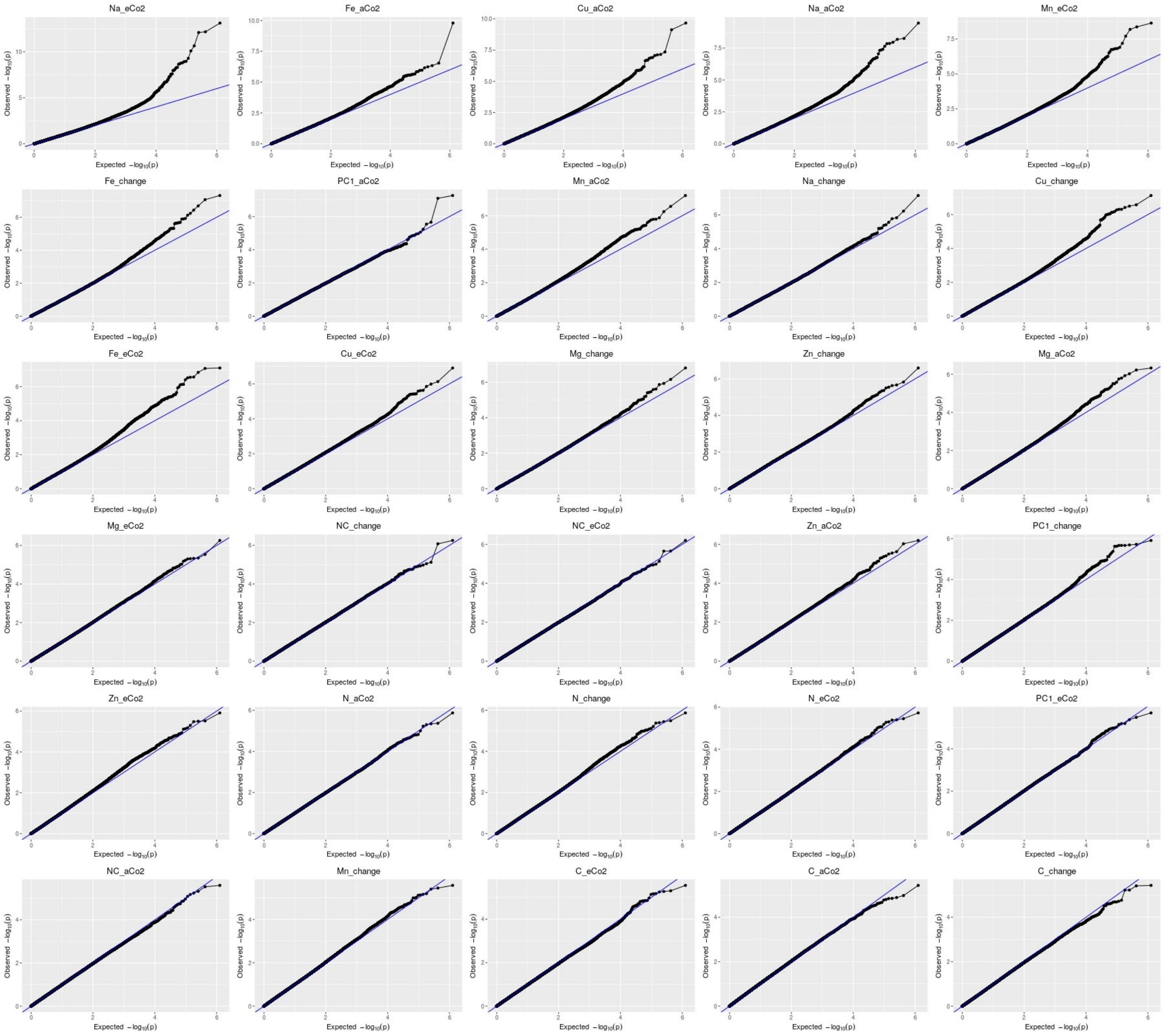


**Supplemental Figure 3:** Qqplots from the GWAs corresponding to data for the level of each mineral under ambient and under elevated $CO_2$, as well as for the percentage of change between ambient and elevated $CO_2$ for each element.

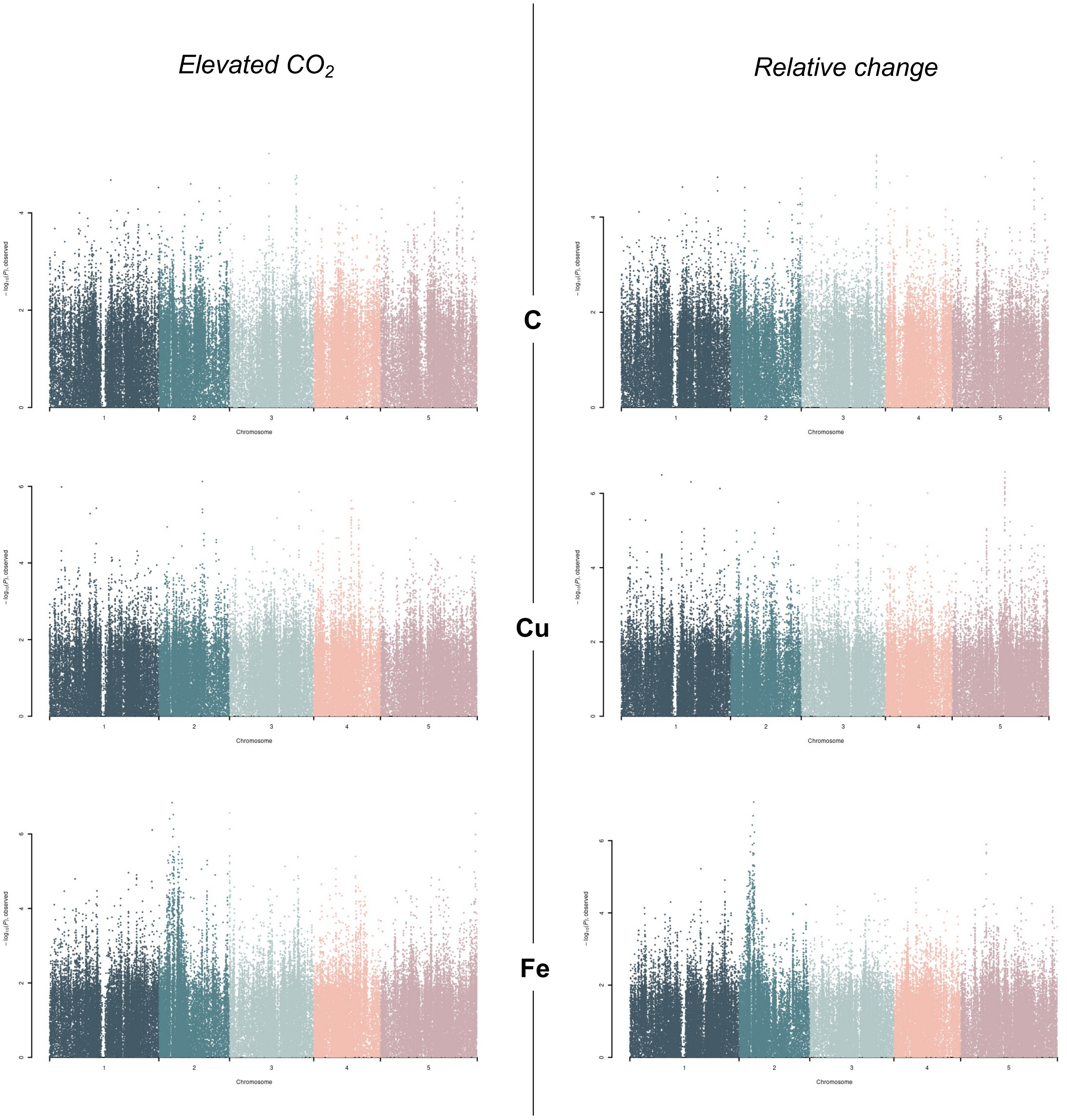


**Supplemental Figure 4:** Manhattan plots made from the GWAs corresponding to data for the level of each mineral under elevated $CO_2$, as well as for the percentage of change between ambient and elevated $CO_2$ for each element.

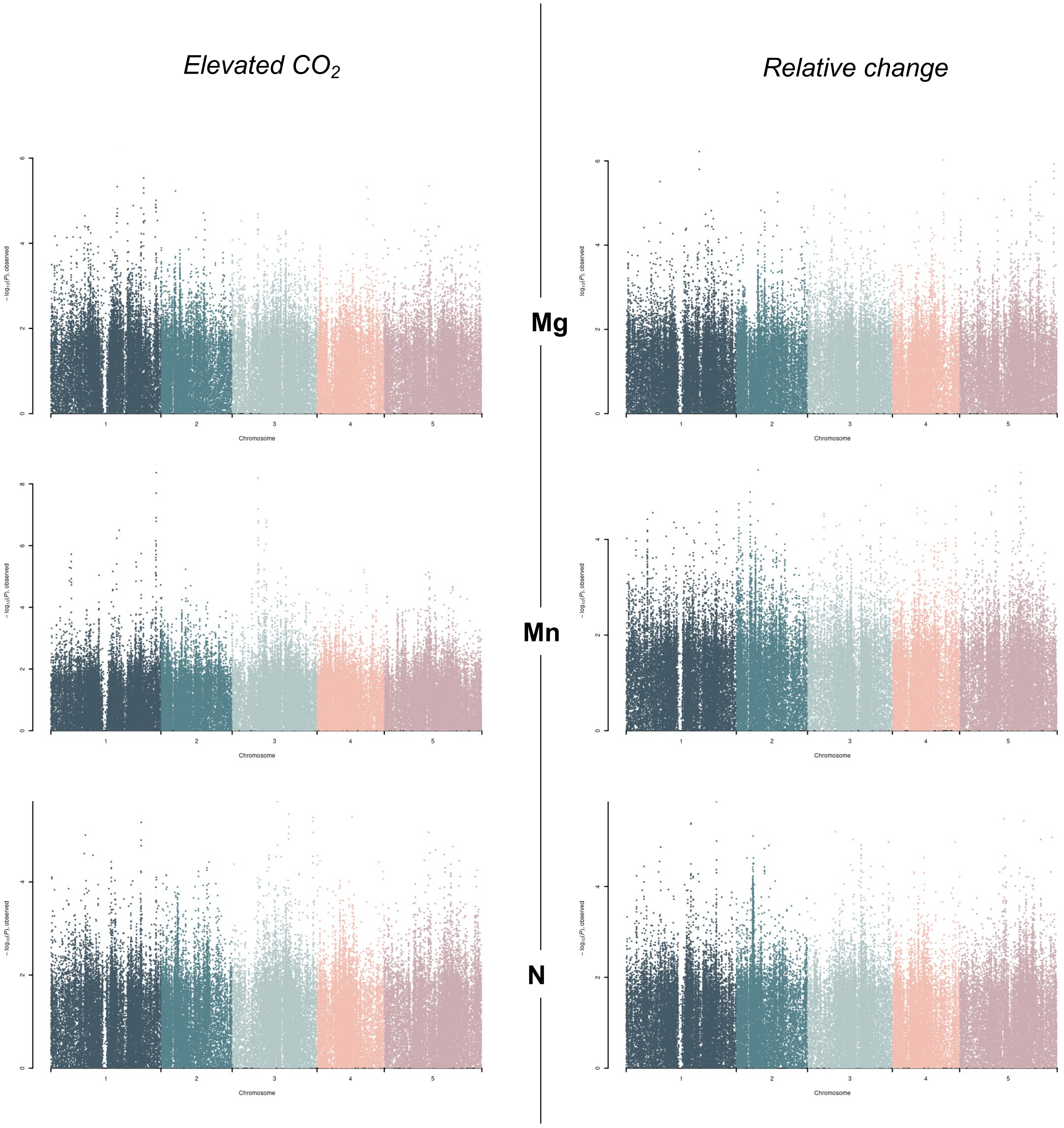


**Supplemental Figure 4 (continued):** Manhattan plots made from the GWAs corresponding to data for the level of each mineral under elevated $CO_2$, as well as for the percentage of change between ambient and elevated $CO_2$ for each element.

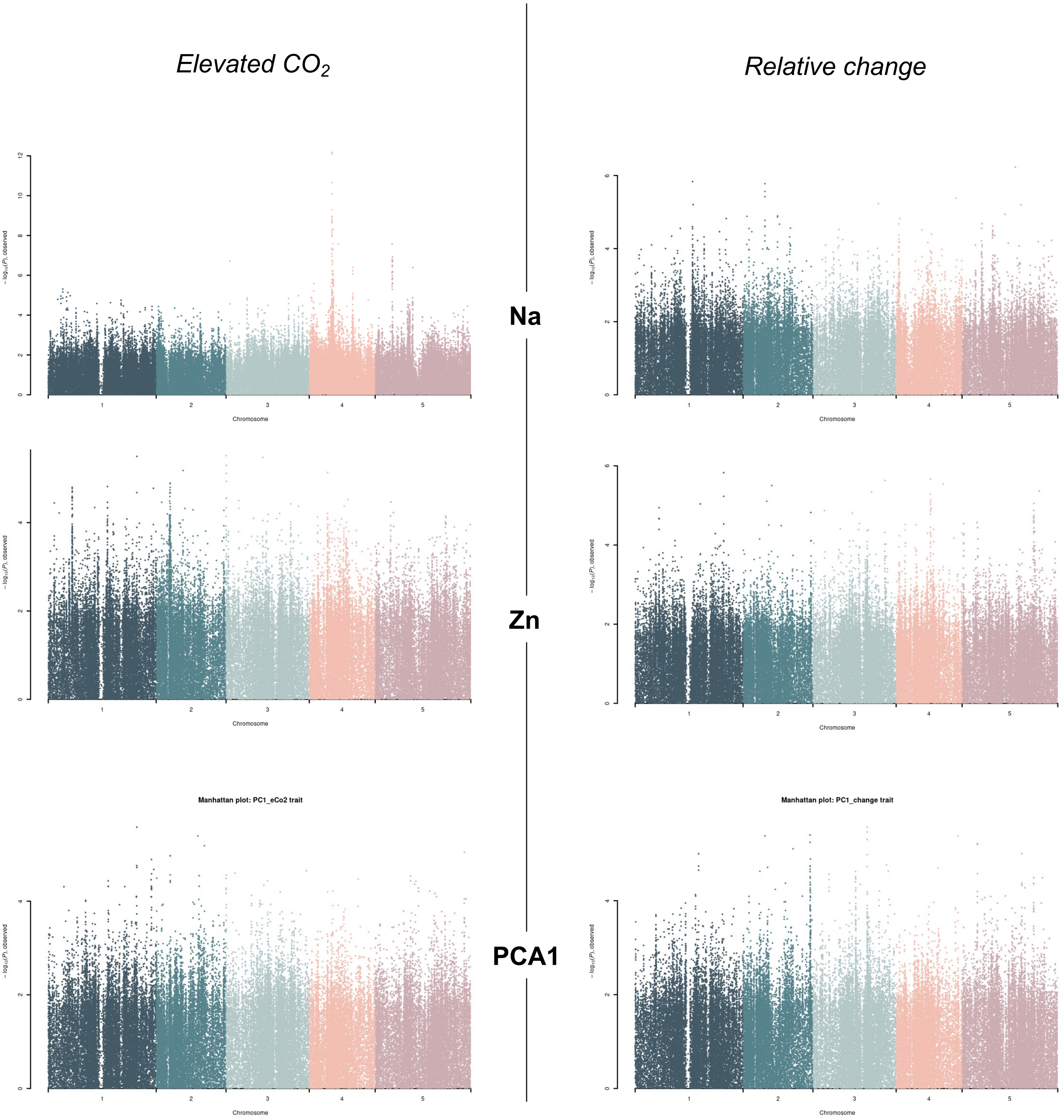


**Supplemental Figure 4 (continued):** Manhattan plots made from the GWAs corresponding to data for the level of each mineral under elevated $CO_2$, as well as for the percentage of change between ambient and elevated $CO_2$ for each element.

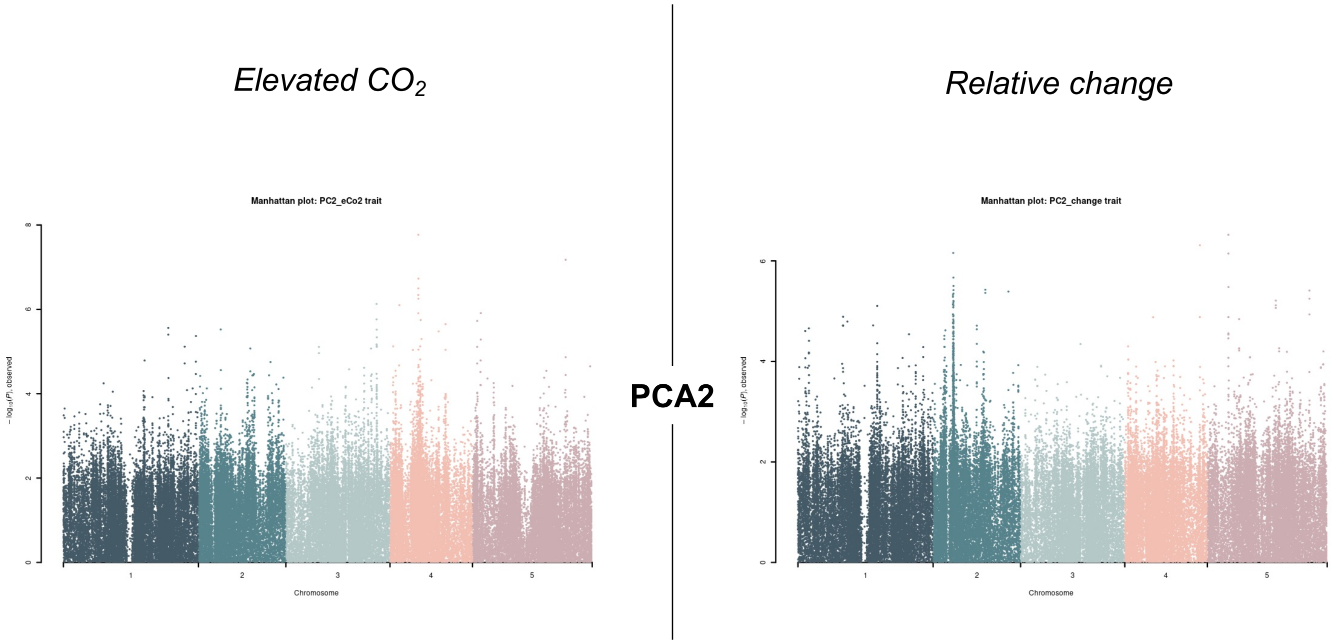


**Supplemental Figure 4 (End):** Manhattan plots made from the GWAs corresponding to data for the level of each mineral under elevated $CO_2$, as well as for the percentage of change between ambient and elevated $CO_2$ for each element.
